# CRGEM: Cellular Reprogramming using mechanism-driven Gene Expression Modulation

**DOI:** 10.1101/2023.05.12.540496

**Authors:** Avani Mahadik, Nitin, Justin Jose, Vivek Singh

## Abstract

**Introduction:** Regenerative medicine promises a cure for currently incurable diseases and pathological conditions. Its central idea is to leverage healthy cells to regenerate diseased cells, tissues or organs through the process of cellular reprogramming. The most common method to achieve this is by modulating the activity of specific transcription factors. However, the large number of protein-coding genes and transcription factors in humans and their complex interactions poses a challenge in identifying the most suitable ones for modulation. Here, we propose a computational workflow that facilitates the prediction of such transcription factors for achieving desired cellular reprogramming, along with highlighting their mechanistic basis in terms of the gene regulatory network of the target cell type.

**Methods:** In this paper, we propose a synergistic workflow that leverages existing computational tools: TransSynW, PAGA and SIGNET, a software – Cytoscape and two databases – TRRUST and UniProt. It uses single-cell transcriptome data of the starting and target cell types as inputs. We demonstrate this workflow by predicting suitable transcriptional modulations for reprogramming of human foreskin fibroblasts to oculomotor neurons.

**Results:** Using the workflow, we hypothesized the core drivers for specific cellular reprogramming along with their functional understanding for experimental applications. The workflow predicted the transcription factors for modulation and provided insight into their differential expression dynamics and influence on the predicted gene regulatory network of the target cells.

**Conclusion:** Our computational workflow helps extract meaningful predictive and mechanistic insights from high-dimensional biological data, which otherwise is difficult to accomplish from individual tools alone. We believe this workflow can help researchers generate mechanistically founded hypotheses for achieving desired cellular reprogramming as a step towards regenerative medicine.

**Highlights:**

- Combine computational tools as workflows to gain predictive and mechanistic insights
- The workflow predicts suitable transcription factors for targeted cellular reprogramming
- Gain insight into the influence of transcriptional modulation on gene regulatory network
- The workflow generates mechanistically founded hypotheses for transcriptional modulation
- Rationalized experimental design for targeted cellular reprogramming for regenerative therapies

## 1. Introduction

Regenerative medicine is an evolving branch of medicine aiming to address diseases without curative treatments [1], [2], [3], [4].The central approach in regenerative medicine is to repair or replace the damaged and diseased cells, tissues and organs with the healthy ones [5]

The first step towards generation of complex tissues and organs is the generation of healthy cells of specific types. This can potentially be achieved by cellular reprogramming through: (1) directed cellular differentiation of pluripotent cells occurring naturally (e.g. embryonic or adult stem cells) [6], [1], or developed artificially (e.g. iPSCs) [7], [8]; and (2) by transdifferentiation of somatic cells directly to the target cells [9].

Cells of a particular type maintain their identity and behavioral properties with the help of dynamic activities of transcription factors (TFs) and their corresponding target genes [10]. In line with this, cellular reprogramming from one type to another is shown to be achieved by modulating the activities of specific transcription factors [11], [12]. However, the challenge lies in identifying the correct TFs for modulation given that there are estimated 19,000 protein-coding genes [13] and ∼1,600 TFs in humans [14]. Therefore, TF modulation for cellular reprogramming is typically based on prior knowledge, empirical intuition or purely trial and error, resulting in low efficiency of reprogramming [8], and tumorigenicity [1] and immune rejection [1], [4] of the target cells in the body.

Technologies such as single cell RNA sequencing (scRNA-seq) provide transcriptome information at single cell resolution [15]. This offers a means to understand the transcriptional signatures of different cell types as well as their intermediate transient cellular states during cellular interconversion [16] and thus, help in identifying TFs whose modulation can bring about the desired cellular reprogramming [17]. In fact, several computational tools exist (e.g. TransSynW [18], CellCartographer [19], NETISCE [20] and scREMOTE [21]) that utilize single cell transcriptomic (sc-transcriptomic) data alone or in combination with complementary biological data, to predict the TF modulation (i.e. gene expression alterations) required for desired cellular reprogramming. However, the hypotheses on TF modulation alone without understanding their molecular basis may be a black box and may have adverse consequences if progressed further towards clinical settings.

To gain holistic understanding of TF modulation, their mechanistic basis and the downstream effects, existing computational tools can be leveraged in the form of analysis workflows. In this study, we propose an analysis workflow called CRGEM, to generate hypotheses on TF modulation for cellular reprogramming and understand their mechanistic basis with respect to the transcriptional regulatory network of the target cell type.

We demonstrate the usefulness of CRGEM on the sample dataset used by Ribeiro et al.[18]. The dataset corresponds to single cell transcriptome data from human foreskin fibroblasts (HFF) and human midbrain development.

## 2. Results and Discussion

We present CRGEM, an integrated analysis workflow consisting of three disparate computational tools: TransSynW[18] is for generating hypotheses on TF modulation for cellular reprogramming of starting cell type to the target cell type; PAGA [22] for understanding the importance of these hypothesized genes in the context of cellular reprogramming process; and SIGNET [23] to understand the TF modulation in the context of gene regulatory network of the target cell type, and gain insight into the mechanistic basis of the TF modulation hypothesis (see Figure 1).

**Figure 1:**
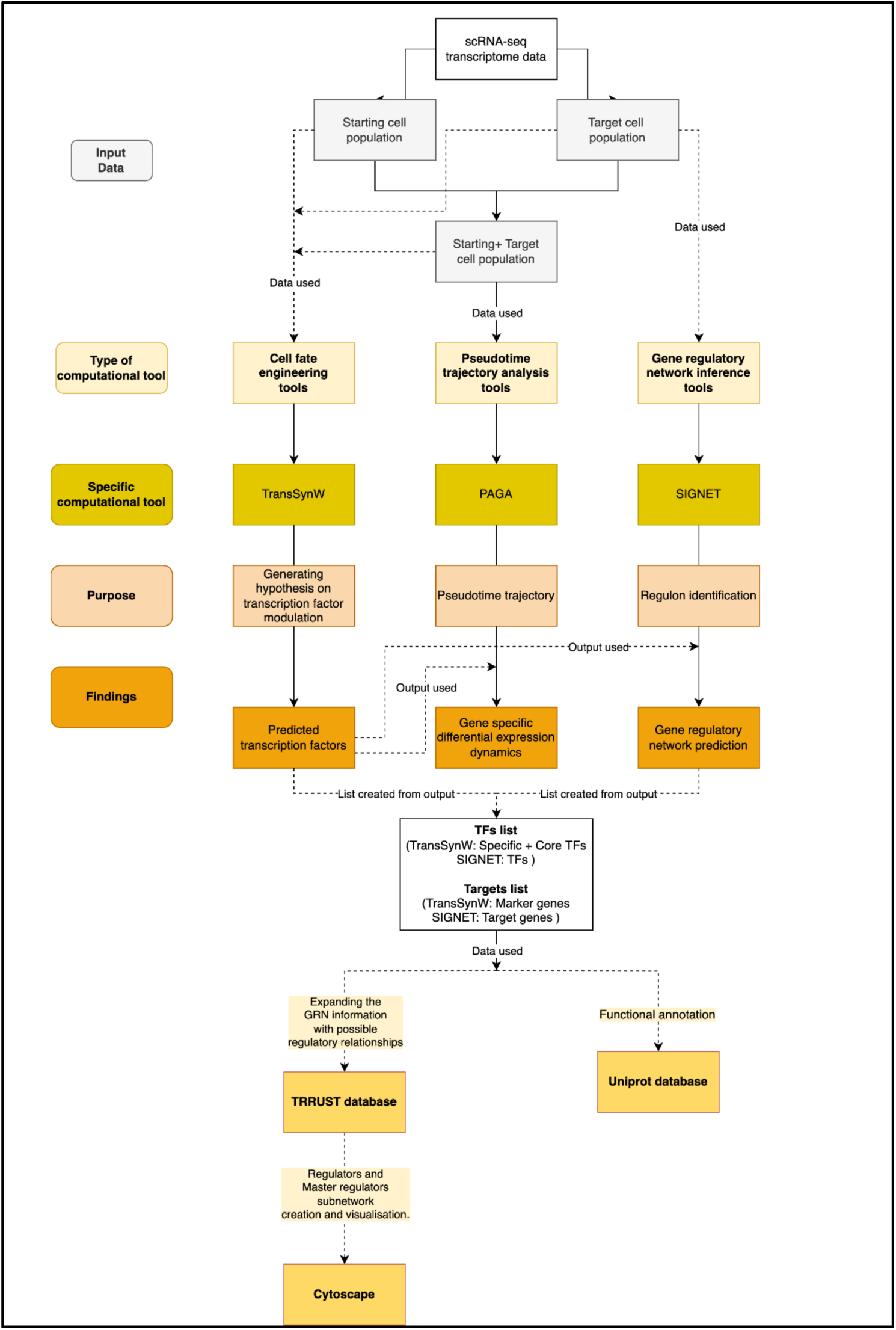
CRGEM – Flow diagram

Furthermore, we used TRRUST [24], a database of human and mouse transcriptional regulatory networks, to augment the list of master regulators and target genes corresponding to the genes predicted by the workflow. Paths from master regulators and TFs to their target genes were identified using Cytoscape [25] (see Methods for detail). This resulted in a gene regulatory network that captures the key players and their effect on transcriptional regulation that bring about the required transcriptional changes required for cellular reprogramming of starting cell type to the target cell type. The functional annotation of these genes were retrieved from the UniProt database.

The proposed analysis workflow is implemented in Python, and the source code is available at: https://github.com/thoughtworks/cell-reprogram-workflow. The schema of the workflow is shown in Figure 1.

We demonstrate this workflow using single cell transcriptomic datasets corresponding to HFF and a midbrain associated cell type called Oculomotor neuron (hOMTN). The datasets were obtained from the NCBI GEO database. HFF cells form the starting cell type, and the goal of the workflow is to generate mechanistically founded hypotheses for converting HFF cells to hOMTN. The workflow generated hypotheses on TF modulation for cellular reprogramming, revealed the differential expression dynamics of the predicted TFs, and their interaction with the predicted gene regulatory network of the target cell type. These results are discussed in detail in the subsequent sections.

CRGEM can be executed in a modular fashion i.e. all the steps can be executed together in a sequence, or the workflow can be executed in stages. The details of these stages are discussed in the Methods section.

### 2.1 Hypothesis generation on transcription factors whose activity modulation can achieve desired cellular reprogramming

We have used TransSynW, a cell fate engineering tool, in the workflow for generation of hypotheses on TF modulations required for desired cellular reprogramming. TransSynW requires single cell transcriptomic data of starting and target cell populations as input.

TransSynW predicts a set of core TFs whose expression modulations are hypothesized to bring about the desired cellular reprogramming. Core TFs are of two types: (1) Pioneer TFs: related to generic functions like chromatin opening required during gene expression alteration, and (2) Specific TFs: specific to the starting and target cell types whose expression modulation is required for desired cellular reprogramming.

TransSynW also predicts marker genes for the target cell type. These genes can be used to assess whether the resulting cells are of the desired target cell type, and thereby, indicate the performance of the cell type reprogramming [18].

In CRGEM, the stage named ‘generate_hypothesis’ performs this task of predicting suitable TF modulations.

In the case of HFF to hOMTN reprogramming, TransSynW predicted 8 core TFs (5 specific factors and 3 pioneer factors), whose activity modulations are hypothesized to bring about the desired cellular reprogramming (see Table 1). The marker genes predicted by TransSynW are shown in Table 2.

**Table 1:** Core transcription factors predicted by TransSynW for reprogramming of HFF to hOMTN.

| Gene | FoldChange | Type of core transcription factor |
| --- | --- | --- |
| PEG3 | 9.493 | specific |
| ***PHOX2B*** | 9.096 | specific |
| CSRNP3 | 7.364 | specific |
| ***PHOX2A*** | 7.299 | specific |
| CXXC4 | 7.132 | specific |
| ***ISL1*** | 9.83 | pioneer |
| *FOXA2* | 4.118 | pioneer |
| *ASCL1* | 3.603 | pioneer |

The genes indicated in bold font correspond to the results presented in the paper. Additionally, genes that are highlighted in bold and italics, not only align with the reported findings but are also experimentally verified.

**Table 2:** Marker genes predicted by TransSynW for reprogramming of HFF to hOMTN. JSD Score: Jensen-Shannon Divergence score

| Gene | JSD score | Subpopulation |
| --- | --- | --- |
| *TUBB2B* | 102.06 | *hOMTN* |
| *DCX* | 102.11 | *hOMTN* |
| *NEFM* | 102.12 | *hOMTN* |
| *CD24* | 102.12 | *hOMTN* |
| *CRMP1* | 102.12 | *hOMTN* |
| *KIF5C* | 102.14 | *hOMTN* |
| *MARCKSL1* | 102.14 | *hOMTN* |
| *MLLT11* | 102.15 | *hOMTN* |
| *ELAVL4* | 102.15 | *hOMTN* |
| *NCAM1* | 102.17 | *hOMTN* |

### 2.2 Gaining insight into the differential expression dynamics of hypothesized transcription factors using pseudotime trajectory analysis

We leveraged PAGA, a pseudotime trajectory inference tool, to understand the relevance of TransSynW predicted transcription factors in the context of cellular reprogramming from HFF to hOMTN.

PAGA uses the starting cell type data (i.e. HFF) as the root (i.e. reference) and attempts to create a pseudotime trajectory representing the degree of reprogramming of each cell to the target cell types. Overlaying the gene expression data of individual genes predicted by TransSynW on this pseudotime trajectory shows the difference in their expression as the cells transition from HFF to the target subtype (hOMTN in this case).

With respect to HFF to hOMTN reprogramming, all the specific *TFs i.e. PEG3, PHOX2B, CSRNP3, PHOX2A and CXXC4* showed distinct change of expression between these cell types (see Figure 2). Among the pioneer genes*, ISL1* showed a clear distinct change of gene expression, while *FOXA2* and *ASCL1* showed distinct expression changes in some samples and not in others (see Figure 3).

**Figure 2:**
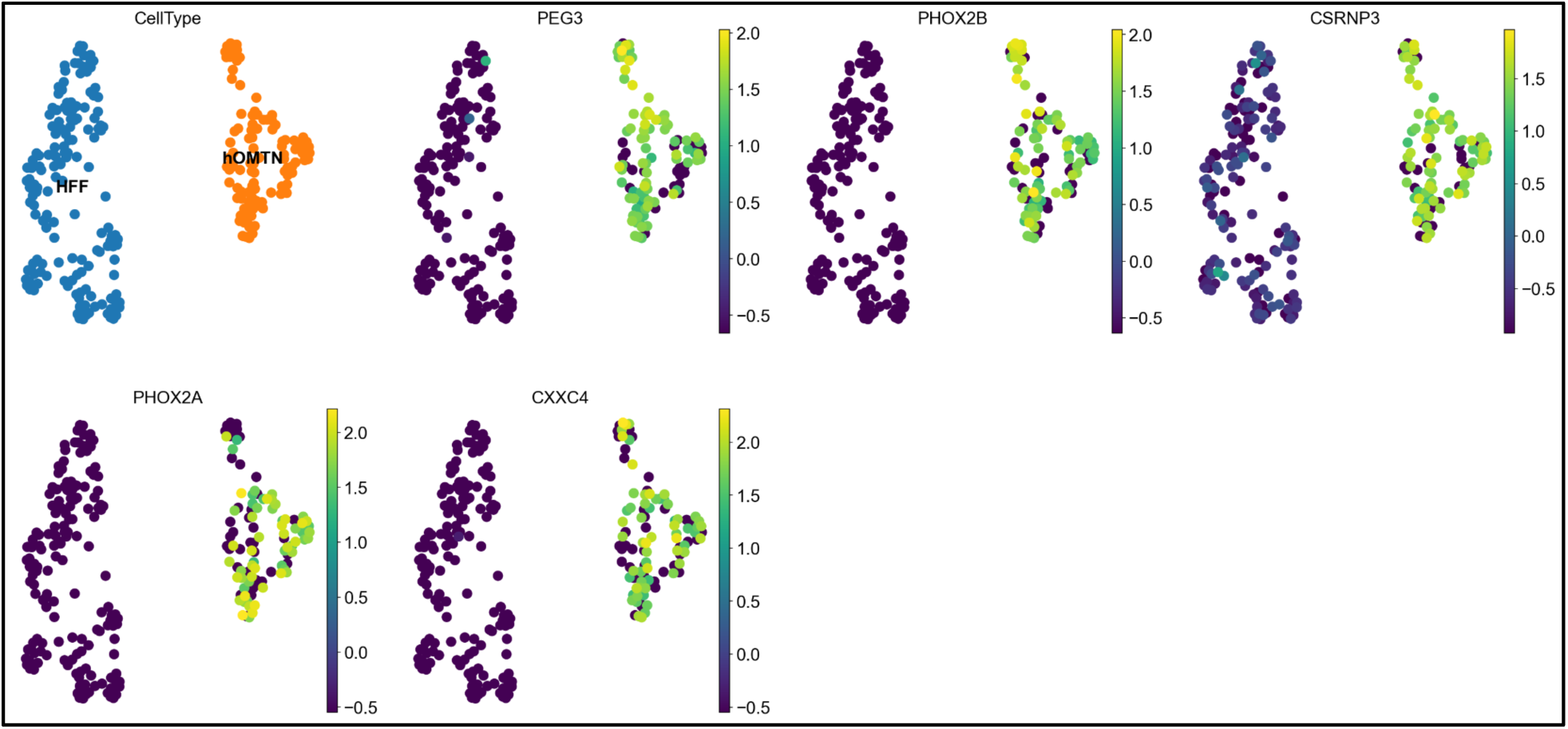
D**i**fferential **expression dynamics of hypothesized specific transcription factors with reference to the pseudotime trajectory.** Clustering of HFF and hOMTN cells is provided for reference; Blue: HFF cells, Orange: hOMTN cells.

**Figure 3:**
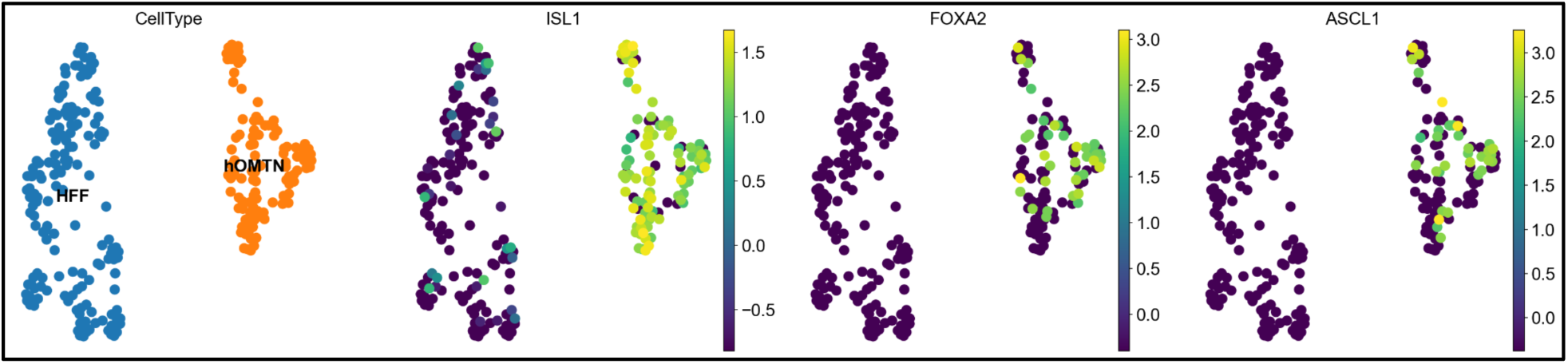
D**i**fferential **expression dynamics of hypothesized pioneer transcription factors with reference to the pseudotime trajectory.** Clustering of HFF and hOMTN cells is provided for reference; Blue: HFF cells, Orange: hOMTN cells.

CRGEM has a stage titled ‘mechanistic_insight’ that integrates TransSynW and PAGA related steps for generating hypotheses and gaining mechanistic insight into the predicted TFs.

### 2.3 Deciphering the mechanistic basis of transcription factors activity modulation

We used SIGNET to predict the gene regulatory networks (GRNs) of the target cell types that highlight the functional transcriptional regulation required for cellular functions (see Table 3). Assessing the TransSynW hypothesized TFs and marker genes in the context of this GRN can reveal the likely molecular basis of their effect on cellular reprogramming. It can also highlight the cascading effect(s) (desired or inadvertent), of TF modulation resulting from regulatory interactions in the GRN.

**Table 3:** Regulons predicted by SIGNET for target cell type, hOMTN.

| TFs | Gene_list |
| --- | --- |
| ATF1 | SYT1, MCFD2 |
| ELF2 | NEDD4L, STX16 |
| FOXA1 | RASSF8, STX16, IRX3, CTBP2 |
| GABPA | CAMK2N1, RPS27 |
| NCALD | RASSF8, KIF5C, PDHA1 |
| POU2F1 | SETD1B, NEDD4L, IRX3 |
| RFX7 | HSPBP1, FOS |
| ZBTB44 | RASSF8, ANKRD11 |
| ZNF300 | RBM34, LMAN1, RPSA, APOO |

Further, TRRUST database was used to identify the corresponding TFs and the target genes of the genes obtained from TransSynW and SIGNET such that indirect interactions and master regulator(s), if any, can be captured.

The resulting disparate regulatory interactions are integrated as a holistic gene regulatory network using Cytoscape and one of its apps called PathLinker [26] (see Figure 4). In particular, PathLinker facilitated computing paths from master regulator(s) to TFs, and TFs to their target genes. Functional annotations of the genes retrieved from the UniProt database and are available in the Supplementary file.

**Figure 4:**
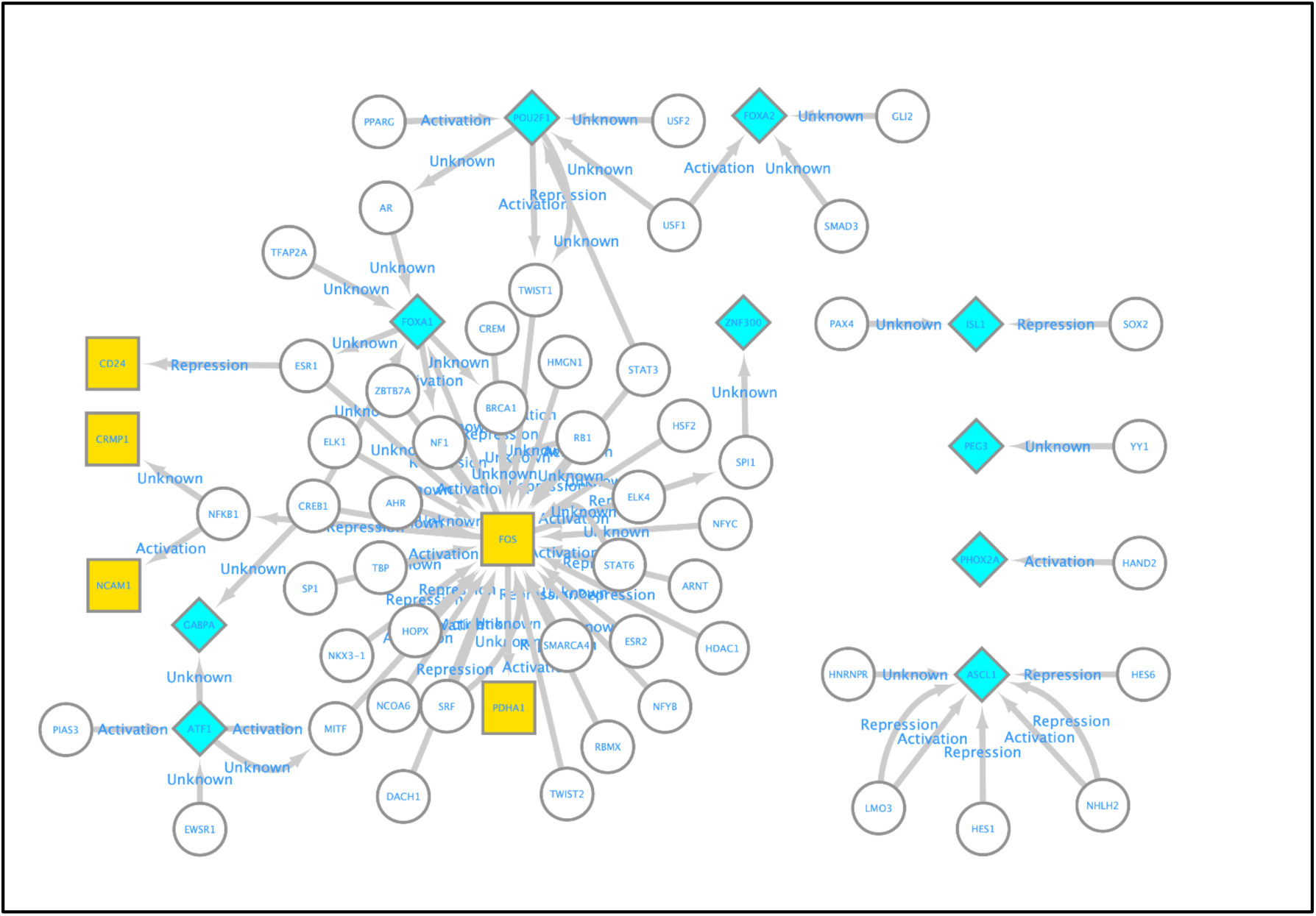
C**o**mprehensive **visualization of gene regulatory network obtained from Cytoscape.** Colour and shape representation: Yellow square shaped nodes represent marker genes and blue diamond shaped nodes represent TFs.

The stage titled ‘grn_inference’ performs the gene regulatory network analysis using SIGNET. The stage trrust_analysis is for extracting the regulators and target genes from the TRRUST database. Gene regulatory network was visualized with a stage named ‘create_network.’ Further stage called ‘functional_annoation’ is used for extracting functional annotations from the UniProt database.

## 3. Discussion

Efficient cellular reprogramming of therapeutically suitable starting cell types (e.g. stem cells, skin fibroblasts) to the desired target cell types (e.g. dopaminergic neurons, cardiomyocytes, pancreatic beta cells [27]) lies at the center of regenerative medicine. We proposed an analysis workflow that leverages three disparate computational tools, two databases and a software to generate not just hypotheses of TF modulation for cellular reprogramming, but also infer their mechanistic basis and functional importance. The workflow is implemented in Python. Users can execute the complete workflow from start to end or can execute different stages provided by the workflow or extend the workflow by adding their additional tools based on specific needs.

Although scientific literature is replete with examples of using multiple tools to address their specific use cases in cellular reprogramming and regenerative medicine, to the best of our knowledge, such defined workflows are not present. We believe that the proposed workflow is reusable and can help researchers formalize new and mechanistically founded hypotheses for achieving desired cellular reprogramming.

It is important to note that the presented workflow is one of the possible workflows, and many more such workflows can be devised to address this or other challenges in the regenerative medicine field. We would like to draw parallels to genomic data analysis workflows [28], [29], [30] that have streamlined such analyses and have contributed greatly in accelerating the progress of the field.

## 4. Conclusion

In this study, we leveraged diverse computational tools together to navigate through high-dimensional transcriptomic data and proposed an analysis workflow, CRGEM, to generate mechanistically founded hypotheses on TF modulation for desired cellular reprogramming. We believe that CRGEM can benefit researchers in devising cellular reprogramming protocols that overcomes the common pitfalls of cellular reprogramming including low efficiency of reprogramming, and the risk of tumorigenicity and immunogenicity in the target cells. Furthermore, we believe integrating already existing resources (computational tools are an example) can in general provide deeper and diverse insights than individual tools. CRGEM is a start and can be further extended with other combinations of tools, prior biological knowledge, etc. based on their suitability to specific use-cases.

## 5. Materials and Method

### 5.1 Single cell transcriptomics-based gene expression data of starting and target cells

The datasets used in the study were obtained from the NCBI GEO database with the following IDs: GSE75748 (for HFF cells) and GSE76381 (for hOMTN). The gene expression values in both the datasets are expressed as counts of detected mRNA molecules from each gene in each cell.

GSE75748 consists of scRNA-seq data profiled from cells derived from human embryonic stem cells. HFF is one of the cell-type clusters in it and was used as control. For our study, data corresponding to HFF cells were obtained i.e. 159 samples out of 1018 samples.

GSE76381 refers to a study of midbrain development, from which hOMTN data is used in this study. The complete data consists of 1978 samples out of which hOMTN has 106 samples.

The common genes from both the datasets are obtained. The GSE75748 dataset has 19096 genes and the GSE76381 dataset has 19530 genes. Common genes between them are 15208. The data corresponding to this common set of genes is used for demonstrating the proposed workflow, CRGEM.

### 5.2 Tools

We need to clone the CRGEM project from GitHub (https://github.com/thoughtworks/cell-reprogram-workflow). Once the project is cloned the user needs to run the setup shell script in order to clone all the dependencies required for working of the tool.

#### 5.2.1 TransSynW

TransSynW was set up locally using code available at: https://git-r3lab.uni.lu/mariana.ribeiro/transsynw The setup file does the task of cloning the repository and placing the code in dependencies directory. The TransSynW code is used directly from the GitLab repository without any changes.

TransSynW requires sc-transcriptomic data of starting and target cells as input. They are provided through a data directory. TransSynW also requires a background population data. By default, all the cell-types data other than the target cells data in the target cell data file is used as the background population data. However, we used the starting cell data as the background population data by appending it to the target cell data file. This was done with the following rationale: (1) to identify TFs that are specifically expressed TFs, starting cells data can serve as a suitable background population; (2) it is relatively easy to find or generate sc-transcriptomic data corresponding to starting and target cell-types, than to also have data for similar cell-types to serve as background population; and (3) the exact requirement for a dataset to qualify as suitable background population is undescribed.

Moreover, TransSynW uses all the genes from starting and target cells for its processing. However, the genes present in one cell-type and missing in the other are then, at times, inferred as specific genes. To avoid such bias, we execute TransSynW using only the common set of genes among starting and target cells.

#### 5.2.2 PAGA

PAGA [22] requires the sc-transcriptomic data in the form of a samples x genes matrix as input. Thus, the target cells data file used for TransSynW, which consists of merged data from start and target cells, is transposed and used as input for PAGA.

Scanpy (https://scanpy.readthedocs.io/en/stable/index.html) was used for PAGA analysis, and the steps are based on its tutorial (https://scanpy-tutorials.readthedocs.io/en/latest/paga-paul15.html). Briefly, dimensionality reduction was performed using Principal Component Analysis, followed by construction of neighborhood graph of the samples. The data was further denoised using diffusion map. Starting cell type (i.e. HFF) data is used as the root for computing the pseudotime trajectory. The gene expression values of TransSynW predicted genes were overlaid on the trajectory to visualize the expression changes of each gene in the context of the trajectory.

#### 5.2.3 SIGNET

SIGNET is implemented as two parts. The first part is in Python, and the corresponding code is obtained from the following link: https://github.com/Lan-lab/SIGNET/blob/main/SIGNET_Tutorial/SIGNET.py. In CRGEM, the setup file retrieves the file and stores it in the designated directory. The second part is in R language. The corresponding R script is written following the steps described in: https://github.com/Lan-lab/SIGNET/blob/main/SIGNET_Tutorial/SIGNET_GRN_prediction.md.

SIGNET [23] requires the target cells sc-transcriptomic data in the form of samples x genes matrix as input. It also requires a list of transcription factors based on the species and gene motifs ranking file which was downloaded from: https://resources.aertslab.org/cistarget/.

This file is not provided in the GitHub repository.

SIGNET predicts the relationship between TF and non-TF genes, and then uses RcisTarget (https://bioconductor.org/packages/release/bioc/html/RcisTarget.html) for motif analysis and thereby filtering out the false positive ones.

#### 5.2.4 TRRUST

TRRUST is a manually curated database of human and mouse transcriptional regulatory networks. For a given gene, it contains information about the TFs regulating it, and the target gene(s) regulated by it. TRRUST database network was downloaded in. tsv format from: https://www.grnpedia.org/trrust/downloadnetwork.php and used in CRGEM. Users can use the provided file or can download its latest version.

TRRUST is used to find the regulators and target genes of the genes obtained from TransSynW and SIGNET. In particular, the TFs obtained from TransSynW and SIGNET are used to retrieve: (1) their regulators i.e. TFs that regulate these TFs, and (2) their target genes. Similarly, the marker genes and target genes obtained from TransSynW and SIGNET respectively are used to retrieve their regulators and the target genes (if any).

#### 5.2.5 Cytoscape

Cytoscape version 3.9.1 was used for all the processing and computations related to the creation of a holistic gene regulatory network in CRGEM. Cytoscape PathLinker app was used to create two sub-networks by computing paths between: (1) TFs from TransSynW and SIGNET, and marker and target genes from TransSynW and SIGNET respectively; (2) Master regulator(s) if any and TFs from TransSynW and SIGNET. These sub-networks are subsequently merged together into one network using Cytoscape. The merged network allows unraveling the master regulator(s) and identifying the paths through which they regulate the TFs. Moreover, it also reveals the paths through which TFs regulate the marker and target genes.

The Python library, py4cytoscape, was used for performing these analyses.

#### 5.2.6 UniProt

UniProt database was programmatically accessed with a Python script, to get the functional annotations of the TFs and target genes. The python script was written following the tutorial https://embl-ebi.cloud.panopto.eu/Panopto/Pages/Viewer.aspx?id=a4e18a75-8b00-43c7-8120-af1f0105b7ed.

## Data availability

We downloaded the gene expression data from Gene Expression Omnibus (GEO) under accession codes GSE75748 and GSE76381. The source code for CRGEM is available at: https://github.com/thoughtworks/cell-reprogram-workflow.

## Abbreviations

HFF: human foreskin fibroblasts
hOMTN: Oculomotor neuron
scRNA-seq: single cell RNA sequencing
sc-transcriptomic: single cell transcriptomic
TFs: Transcription Factors

## Acknowledgment

The authors would like to thank Dr. Mariana Ribeiro for addressing our TransSynW related queries. We would also like to thank Harshal Hayatnagarkar, Divye Singh and Anuja Chaitanya for valuable discussions and helping us in troubleshooting technical issues with installation and usage of the tools.

